# An Arrayed Transposon Library of *Ruegeria pomeroyi* DSS-3

**DOI:** 10.1101/2022.09.11.507510

**Authors:** Catalina Mejia, Lidimarie Trujillo Rodriguez, Ravin Poudel, Adam Ellington, Adam R. Rivers, Christopher R. Reisch

## Abstract

The ability to construct defined genetic mutations in many bacteria is difficult and limited. Transposon mutagenesis is often highly efficient, but is not site specific, thus selections are often needed to identify mutants of interest. The construction of arrayed mutant libraries would help to fill this need, though these libraries are costly and time consuming. To enable easier construction of arrayed libraries we developed a workflow and methodology using a hierarchical barcoding scheme to identify mutants within a multiwell plate. We applied this method to the marine Alphaproteobacterium *Ruegeria pomeroyi* DSS-3 and created a library with over 2,800 disrupted genes.

## Introduction

Heterotrophic marine bacteria shape global biogeochemical cycles and drive nutrient cycling in the surface oceans (Karl & Church, 2014). While there has been much progress over the last two decades in understanding the genetic underpinnings of these cycles, many gaps remain. The explosion of omics data collection over the past two decades has provided extensive data at the community level, enabling hypothesis generation (Faure et al., 2021). More generally, the number of genes with unknown function in bacteria remains high. Even with the most recent computational tools, about 30% of microbial genes have an unknown function (Vanni et al., 2022).

Transposon mutant libraries and next generation sequencing have enabled systems level investigations of bacterial genetics and physiology. Barcoded transposons and the associated Bar-seq workflow has further simplified these genome level analyses (Wetmore et al., 2015). In Bar-seq, each transposon possesses a 20-bp random barcode. After an initial Tn-Seq to identify the genomic location and barcode sequence of the transposon, only amplification of the 20-bp barcode is needed for subsequent experiments, greatly speeding and simplifying the workflow. In fact, this method was used to provide phenotypic data for 11,779 proteins with unknown function (Price et al., 2018).

Starting with a barcoded library, Shiver et al. described a technique to construct an ordered library of mutants in a strict anaerobic bacterium (Shiver et al., 2021). This technique used a grouped pooling approach, where the location of each mutant in the plate is deconvoluted computationally. Similarly, we describe a method for constructing an arrayed transposon mutant library that relies on the 20-bp barcode for identification. However, our method uses a hierarchical barcoding scheme to identify the row, column, and plate where each mutant originated. A single PCR reaction is performed with each mutant arrayed in a 384-well plate as template. A series of 16 and 24 primers that correspond to each row and column of a 384-well plate. The products from each plate are then pooled and amplified with Illumina indices. Each amplicon thus contains the information to identify the 20-bp barcode, column and row position, and plate from which the barcode originated, in a technique dubbed PlateSeq. We applied PlateSeq to the marine Alphaproteobacterium *Ruegeria pomeroyi* DSS-3 and obtained arrayed mutants for over 2800 genes, demonstrating feasibility of the methods and providing a useful set of genetic mutants for research. An overview and flow chart of the methods used for the Tn-Seq and subsequent arraying and plate mapping is given in Figure 1.

**Figure 1.**
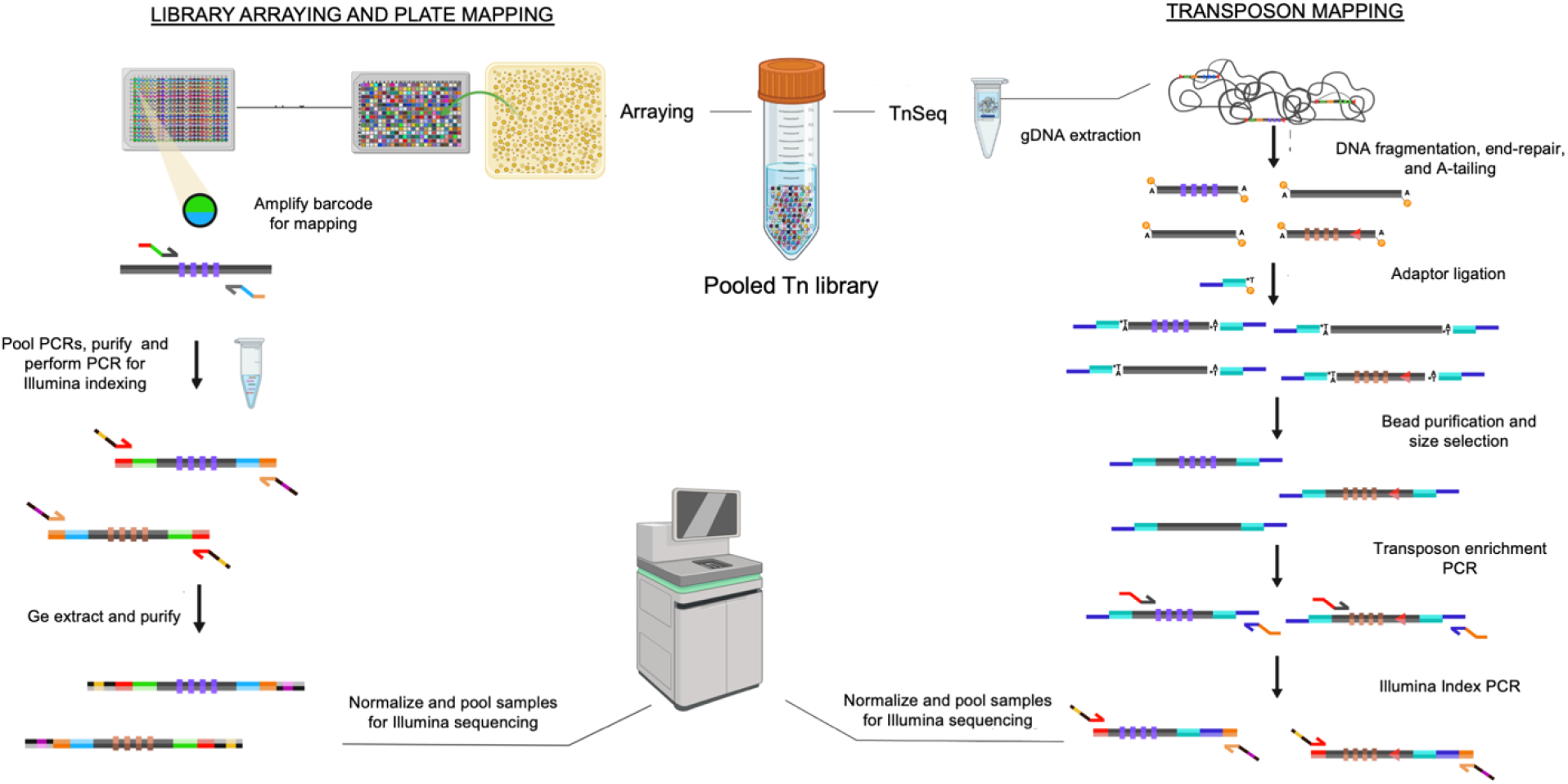
Overview of TnSeq and PlateSeq workflow. After transposon library construction, TnSeq and BarSeq are depicted on the right side of the diagram. Preparation of the arrayed libraries and subsequent PlateSeq methods are depicted the left-side of the diagram. Figure created with BioRender.com..

## Materials and methods

### Media

1. ½ YTSS broth is prepared by adding 2 g of yeast extract, 1.25 g of tryptone, 20 g of sea salts to 1 L of deionized water (dH2O).
2. Luria-Bertani broth (LB) is prepared by adding 10 g of tryptone, 5 g yeast extract, and 10 g NaCl.
3. 100 mg L^-1^ kanamycin (kan) in dH2O
4. 150 mM Diaminopimelic acid (DAP)
5. For solid media, add 15 g/L agar.
6. 384-well plates with 80 μL of ½ YTSS with 100 mg L^-1^ kan
7. 384-well plates with 60 μL of ½ YTSS with 100 mg L^-1^ kan
8. Sterile 80% glycerol

### Bacterial strains

1. *R. pomeroyi* is stored in 20% glycerol at −80 C and grown on ½ YTSS medium.
2. *E. coli* WM3064 with barcoded pkmw7 is stored in 20% glycerol stock at −80 C. and grown on LB with 100 mg L^-1^ kan
3. *R. pomeroyi* Tn-5 library is stored in 20% glycerol stock at −80 C and grown on ½ YTSS with solid media with on 100 mg L-1 kan.

### Reagents

1. 0.25% xylene cyanole dye is prepared by adding 0.0125 g of xylene cyanole and 3.53 mL 85% glycerol to 1.47 mL dH2O and filtering through a 0.2 μM filter.
2. 25 mM Deoxynucleotide (dNTP) Solution Mix
3. 5x OneTaq Standard Reaction Buffer (New England Biolabs)
4. OneTaq DNA polymerase (New England Biolabs)
5. Q5 High-Fidelity 2X Master Mix (New England Biolabs)
6. 1X OneTaq Master Mix is prepared according to manufacturer protocol. 1X OneTaq Master Mix
7. DNA gel extraction kit
8. PCR purification kit
9. NEBNext Ultra II FS DNA Library Prep with Sample Purification Beads (NEB #E6177)
10. 80% Ethanol (freshly prepared)
11. Nuclease-free water
12. 0.1X TE (pH 8.0)
13. Primers

a. TA_Adaptor_Top - /5Phos/CTCACCGCTCTTGTAGCTGTCTCTTATACACATCTCCGAGCCCACGAGAC
b. TA_Adaptor_Bottom - CTACAAGAGCGGTGAGT
c. 2815: 5’-CCTACACGACGCTCTTCCGATCTGTCTCGTGGGCTCGGAGATGTGTATAAG-3’
d. 2814: 5’-GAGTTCAGACGTGTGCTCTTCCGATCTTCACTCACCGCCCTGCAGGGATGTCCACGAGGTCTC-3’
14. BSUp and BSDn Primers Listed in Supplementary table 1
15. Unique Dual Index (UDI) Illumina Adapters

| Component | 1 mL stock | Final Concentration |
| --- | --- | --- |
| **5X OneTaq Standard Reaction Buffer** | 200 µL | 1X |
| **25 mM dNTPs** | 8 µL | 200 µM |
| **OneTaq DNA Polymerase** | 2.5 u |  |
| **0.25 % xylene cyanole dye** | 20 µL | 0.005 % |
| **Nuclease-free water** | 769.5 µL | - |

### Consumables

1. 384-well polypropylene plates
2. 384-pin replicator
3. 384-well polypropylene PCR plates
4. Ultracold compatible aluminum seals
5. Polypropylene PCR microplate seals

### Equipment

1. Opentrons OT-2 liquid handler
2. Opentrons 8-Channel Electronic Pipette P20
3. Multidrop 384 well plate dispenser
4. Thermo Scientific E1-ClipTip Electronic Multichannel Pipettes 16-channel
5. Microplate film sealing roller
6. Genetix Qpix2 colony-picker
7. Thermocycler

## 1. LIBRARY CONSTRUCTION

We constructed the barcoded transposon library by conjugating *R. pomeroyi* with *E. coli* WM3064 containing the pKMW7 Tn5 library (strain APA766) (Wetmore et al., 2015). A filter mating protocol was used here (Stoudenmire et al., 2019), though other methods may be suitable for different strains.

### Filter mating

1. From isolated colonies, grow cultures of *E. coli* donor in 5 mL LB broth with 100 mg L^-1^ kan supplemented with DAP (0.3 mM) at 37° C and *R. pomeroyi* recipient in 5 mL ½ YTSS at 30° C for approximately 16 hours.
2. Dilute cultures 100-fold in 5 mL of respective broth with antibiotic and supplement and incubate in respective incubation temperature until they have reached mid-log phase.
3. Centrifuge 500 μL of *E. coli* donor with ½ YTSS at 4,000 x g for 2 min to remove antibiotic, discard supernatant and wash twice.
4. Resuspend in 500 μL of ½ YTSS.
5. Combine 500 μL of *R. pomeroyi* mid-log culture with the 500 μL of the washed *E. coli* donor.
6. Run mating mix through membrane filter and wash with 2-3 times that volume of ½ YTSS supplemented with DAP (0.3 mM).
7. Remove membrane filter from filter flask and place in ½ YTSS. Vortex to remove the cell from the filter and discard the filter.
8. Centrifuge mix at 4,000 x g for 2 min, resuspend in 1 mL of 1/2 YTSS and wash twice to remove DAP.
9. Plate 500 μL (or a volume that obtains isolated colonies) on ½ YTSS with kan.
10. Incubate plate at 30° C for approximately 48 hours.
11. Collect transconjugant colonies by scraping the plate or washing with ½ YTSS broth. Mix with 50 % glycerol and store at −80° C.

## 2. TRANSPOSON MAPPING PREPARATION

### Genomic DNA extraction of library

Genomic DNA can be extracted with the user’s method of choice. We prefer the method below because it is fast and yields high purity DNA.

1. Pellet 1 mL of the transposon library by centrifugation at about 5,000 x g for 5 min and decant the supernatant.
2. Resuspend the pellet in sterile dH2O for a total volume of 500 μL.
3. Add 500 μL of phenol/chloroform/isoamyl alcohol (25:24:1) and pipette to mix.
4. Centrifuge mixture at 12,000 x g for 5 min.
5. Transfer the aqueous layer to a clean 1.5 mL tube, avoiding the white film and organic layer.
6. Purify using a column-based purification kit and elute in 30 μL of elution buffer.
7. Quantify genomic DNA (gDNA) by QuBit and ensure genomic DNA is clean-with 260/230 and 260/280 being greater than 1.8. Store gDNA at −20 ° C.

### Transposon sequencing library (TnSeq) preparation

Our group adapted a transposon insertion sequencing library preparation *protocol*(*Transposon Insertion Sequencing* (*Tn-Seq*) *Library Preparation Protocol - Includes UMI for PCR Duplicate Removal*, n.d.) and scaled it down by half the total reaction amount. Instead of inserting UMIs into sequenced regions, our 20-bp inline barcodes identifies the transposon insertion before subsequent PCR amplification… Our top adapotor’s top and bottom primer sequences also differ in that…

#### Part 1: DNA Fragmentation, end-repair, and adaptor ligation

Before beginning:

1. Fully thaw and quickly vortex all reagents to mix; briefly spin-down reagents.
2. Once thawed and mixed, keep all reagents on ice.

Adaptor annealing:

1. To anneal the adaptor, mix 15 μL adaptor top F Primer (100 μM), 15 μL 2730 adaptor bottom R Primer (100 μM), and 70 μL nuclease-free H2O in a 0.2-mL PCR tube.
2. Run the following annealing program in thermocycler.

| Step | Temperature | Incubation time | Cycles |
| --- | --- | --- | --- |
| **Annealing** | 98°C | 4 min. | 1 |
|  | 98°C -0.1°C/cycle | 1 sec | 940 |
| **Hold** | 4°C | ∞ | 1 |

Enzymatic fragmentation, end-repair, and ligation:

1. Aliquot 50 ng of library gDNA into a 0.2mL PCR tube and use 1X TE to bring total volume to 13 μL.
2. To fragment DNA add 3.5 μL Ultra II FS Buffer and 1 μL Ultra II FS Enzyme Mix. Pipette up and down several times to mix.
3. Run the following program in a thermocycler with heated lid set to 75°C:

| Step | Temperature | Incubation time | Cycles |
| --- | --- | --- | --- |
| **Fragmentation** | 37°C | 10 min. | 1 |
| **Heat inactivation** | 65°C | 30 min. | 1 |
| **Hold** | 4°C | ∞ | 1 |

Sample can be stored at −20 °C. However, this is not recommended because yield is often decreased.

1. Add 15 μL Ultra II Ligation Mix, 0.5 μL Ligation Enhancer, and 1.25 μL Palani adaptor to reaction, mix well.
2. Run the following NEBNext Ligation program in thermocycler with heated lid off.

| Step | Temperature | Incubation time | Cycles |
| --- | --- | --- | --- |
| **Ligation** | 20°C | 15 min. | 1 |

DNA Cleanup and Size Selection with Sample Purification Beads:

1. Transfer all 34.25 μL of adaptor-ligated reaction to a 1.5mL microcentrifuge tube and add 15.75 μL 0.1x TE (dilute provided 1x TE 1:10) to bring volume to 50 μL. Pipette to mix.
2. Briefly vortex NEBNext Sample Purification Beads and add 15 μL to sample. Pipette to mix.
3. Incubate at room temperature for 5 min, briefly spin down, then place in magnetic tube rack.
4. Once solution becomes clear (about 3-5 min), transfer supernatant (65 μL) to a new 1.5 mL microcentrifuge tube.
5. Add an additional 7.5 μL magnetic beads and pipette to mix. Incubate at room temperature for 5 min, briefly spin down, then place on magnetic tube rack.
6. Once the solution becomes clear (about 3-5 min), remove and discard supernatant.
7. Add 200 μL 80% EtOH, avoiding disturbing the bead pellet. Let sit for 30 sec, then remove and discard EtOH.
8. Repeat step (7) once more.
9. Open tube cap to dry the bead pellet at room temperature for up to 5 min. Carefully aspirate by pipette any remaining droplets if present, without disturbing bead pellet. Bead pellet is dry when dark brown, glossy, and no remaining droplets are visible. Do not over dry- if bead pellet becomes light brown and cracked, sample recovery will fail.
10. Remove from magnetic rack and add 10 μL EB (warmed to 55°C). Pipette to mix and briefly spin down. Incubate for 2 min at room temperature, then place back on magnetic rack.
11. Once clear, transfer supernatant to new 1.5 mL microcentrifuge tube. Proceed to Part 2: Transposon Enrichment PCR or store sample at −20° C

#### Part 2: Transposon Enrichment PCR

1. To amplify the transposon junction, set up the following reaction in a 0.2 mL PCR tube.

| Component | 25 μL reaction | Final concentration |
| --- | --- | --- |
| **Ultra II Q5 Master Mix** | 12.5 μL | 1X |
| **10 μM Forward Primer: 2814** | 2.5 μL | 1 μM |
| **10 μM Reverse Primer: 2815** | 2.5 μL | 1 μM |
| **Fragmented DNA** | 7.5 μL | variable |

1. Run the following NEBNext Enrichment PCR program in thermocycler.

- Run the following NEBNext Enrichment PCR program in thermocycler.
| Step | Temperature | Time | Cycles |
| --- | --- | --- | --- |
| **Initial denaturation** | 98° C | 30 sec | 1 |
| **Denaturation** | 98° C | 30 sec | 15 |
| **Annealing** | 72° C | 15 sec |  |
| **Extension** | 72° C | 15 sec |  |
| **Hold** | 4° C | ∞ | 1 |

Perform 1.2x Magnetic Bead DNA Cleanup as follows.

1. Transfer all 25 μL of enrichment PCR to a 1.5mL microcentrifuge tube and add 25 μL 0.1x TE (dilute provided 1x TE 1:10) to bring volume to 50 μL. Pipette to mix.
2. Briefly vortex NEBNext Sample Purification Beads and add 60 μL to sample. Pipette to mix.
3. Incubate at RT for 5 min, briefly spin down, then place in magnetic tube rack.
4. Once the solution becomes clear (about 3-5 min), remove and discard supernatant.
5. Add 200 μL 80% EtOH, avoiding disturbing the bead pellet. Let sit for 30 sec., then remove and discard EtOH.
6. Repeat step (5) once more.
7. Open tube cap to dry the bead pellet at room temperature for up to 5 min. Carefully aspirate by pipette any remaining droplets if present, without disturbing bead pellet. Bead pellet is dry when dark brown, glossy, and no remaining droplets are visible. Do not over dry- if bead pellet becomes light brown and cracked, sample recovery will fail.
8. Remove from magnetic rack and add 10 μL Buffer EB (warmed to 55°C). Pipette to mix and briefly spin down. Incubate for 2 min at room temperature, then place back on magnetic rack.
9. Once clear, transfer supernatant to new 1.5 mL microcentrifuge tube.
10. Quantify cleaned-up enriched DNA by Qubit, using 1 μL. Record concentration. Proceed to Part 3 or store sample at −20 °C.

#### Part 3: Indexing PCR for Illumina Sequencing

1. To anneal Unique Dual Indexes to enriched DNA, set up the following reaction in 0.2mL PCR tube:
2. Run the following NEBNext Index PCR program in the thermocycler.
3. Perform 1.2x Magnetic Bead DNA Cleanup as follows:

1. Transfer all 25 μL of indexing PCR to a 1.5 mL microcentrifuge tube and add 25 μL 0.1x TE (dilute provided 1x TE 1:10) to bring volume to 50 μL.
2. Briefly vortex NEBNext Sample Purification Beads and add 60 μL to sample.
3. Incubate at room temperature for 5 min, then place in magnetic tube rack.
4. Once the solution becomes clear (about 3-5 min), remove and discard supernatant.
5. Add 200 μL 80% EtOH, let sit for 30 sec, then remove and discard EtOH.
6. Repeat step (5) once more.
7. Open tube cap to dry the bead pellet at room temperature for up to 5 min. Bead pellet is dry when no remaining droplets surround it. It should be dark brown and glossy. Do not over dry. If it becomes light brown and cracked, sample recovery will fail
8. Remove from magnetic rack and add 10 μL EB (warmed to 55°C), incubate for 2 min at room temperature, then place back on magnetic rack.
9. Once clear, transfer supernatant to new 1.5 mL microcentrifuge tube.
4. Quantify cleaned-up indexed DNA by Qubit. Concentration should be higher than the recorded concentration of cleaned-up enriched DNA that was obtained in Part 2, step 10 of this protocol.
5. Assess the size and quality of cleaned-up indexed DNA by Bioanalyzer before sending for Illumina sequencing.

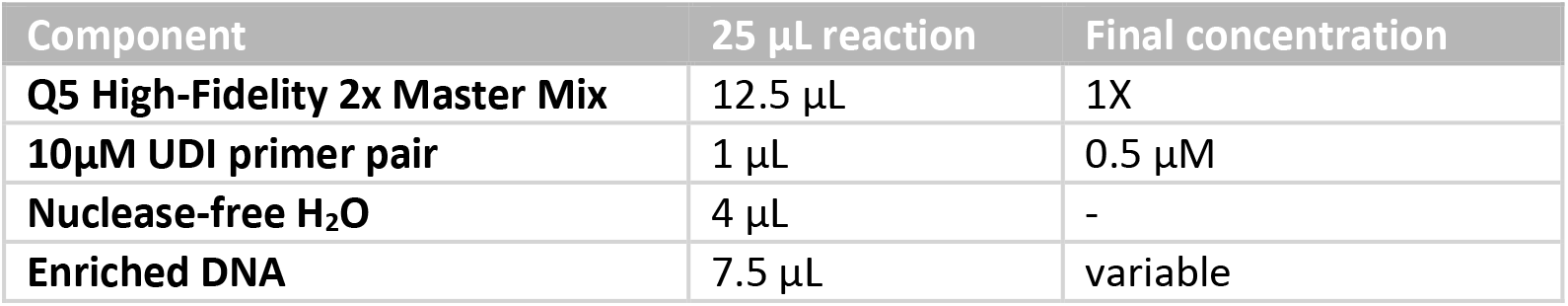

2. Run the following NEBNext Index PCR program in the thermocycler.
| Step | Temperature | Time | Cycles |
| --- | --- | --- | --- |
| **Initial denaturation** | 98 °C | 30 sec. | 1 |
| **Denaturation** | 98 °C | 30 sec. | 7 |
| **Annealing** | 72°C | 15 sec. |  |
| **Extension** | 72°C | 1 sec. |  |
| **Hold** | 4 °C | ∞ | 1 |

## 3. LIBRARY ARRAYING AND PLATE MAPPING (Figure 1)

1. Spread the transposon mutant library onto agar plates so that isolated colonies are obtained after 2-3 days of incubation at 30° C.
2. Inoculate each well with a single colony from agar plate and incubate at 30 ° C for 2-3 days until growth is visible in wells. This original microplate will serve as inoculum for the replicate microplate and for the subsequent PCR protocol.
3. Replicate the 384-well culture plate into a plate with 60 μL of ½ YTSS with 100 mg L^-1^ kan using a 384-pin replicator and incubate at 30° C for 2-3 days.
4. Seal the original microplate with aluminum seal and store in - 80° C.
5. Add 20 μL of 70 % glycerol solution with a multidrop reagent dispenser to each well of the replicate microplates.
6. Seal replicate microplate with aluminum seal and invert 3-5 times to mix glycerol with culture in each well and store at −80° C. The replicate microplate serves as long-term storage of bacterial glycerol cultures.

### Primer plate preparation and PCR 1

1. In a deep 384-well plate, combine the 16 BSUp and 24 BSDown primers so that a unique primer pair is in each 384-well at a final concentration of 0.25 μM.
2. Distribute 5-10 μL of the primers into 384-well PCR plates using the Opentrons liquid handler.
3. Dehydrate the primer aliquots by incubating at 72° C until liquid is no longer visible. The aliquoted plates can be stored with desiccation until usage.
4. Aliquot 5-10 μL of the 1X OneTaq master mix into each well of the 384-well primer PCR plate. Keep on ice or store at 4 ° C until needed.
5. Thaw the stored 384-well culture plates from the −80° C and then briefly centrifuge to remove culture from the aluminum seal.
6. Remove the aluminum seal slowly to avoid cross contamination by splatter.
7. Sterilize the 384-pin replicator with 70% isopropyl alcohol and flame to evaporate alcohol. Allow replicator to cool and dry before proceeding to the next step.
8. Stamp replicator into the 384-well culture plate and lightly mix within wells to pick up settled cells.
9. Stamp replicator into the 384-well primer PCR plate containing 5 μL of 1X OneTaq. Ensure that pins reach the bottom of each well in the PCR microplate.
10. Repeat the above steps to make a duplicate PCR microplate.
11. Seal the duplicate plates with polypropylene PCR plate seals.
12. Briefly centrifuge the plates, so that contents reach the bottom of the wells.
13. Place PCR plates in thermocycler and run the following program:
14. Using a multi-channel pipette or the Opentrons liquid handler, transfer 5 μL of the completed PCR in each well to a 96-well plate. Collect all the reactions that correspond to one 384-well PCR plate in a 1.5-2 mL microcentrifuge tube.
15. Continue this process until all “Step 1” PCRs have been consolidated from one 384-well PCR plate to one microcentrifuge tube accordingly and store at –20° C.
16. Resolve 5 μL of the unpurified pooled Step 1 PCR on a 2% agarose gel to ensure size of about 120 bp.
17. Purify the consolidated pool using a spin column purification kit.

| Step | Temperature | Time | Cycles |
| --- | --- | --- | --- |
| **Initial denaturation** | 94° C | 5 min. | 1 |
| **Denaturation** | 94° C | 30 sec. | 35 |
| **Annealing/Extension** | 68° C | 30 sec. |  |
| **Final Extension** | 68° C | 2 min. | 1 |
| **Hold** | 4° C | ∞ | |

### PCR 2

1. Amplify 300-600 ng of the “Step 1” product to add unique dual indexes and Illumina adapters with Q5 polymerase using the following program.
2. Run “Step 2” PCR product on a 2% agarose gel. The product size should be about 250 bp.
3. Excise the gel fragment and store in a 1.5 mL microcentrifuge tube.
4. Gel purify using a column purification kit.
5. Quantify purified DNA by Qubit.
6. For Illumina sequencing, pool samples at equal concentrations.

| Step | Temperature | Time | Cycles |
| --- | --- | --- | --- |
| **Initial denaturation** | 98° C | 5 min. | 1 |
| **Denaturation** | 98° C | 30 sec. | 10 |
| **Annealing/Extension** | 72° C | 30 sec. |  |
| **Final Extension** | 72° C | 2 min. | 1 |
| **Hold** | 4° C | ∞ |  |

## 4. COMPUTATIONAL MAPPING

Illumina paired end .fastq reads are processed using the FEBA program described previously (Wetmore et al., 2015), the code is available in bitbucket (https://bitbucket.org/berkeleylab/feba/src). The FEBA workflow provides a table output with the barcodes and genomic insertion locations (Tnpool.txt converted to a .csv file). These tables are then used by our in-house programs that identify the plate location for each barcode. These programs are made up of the KEIO.py script, keio.R script and the process.R script. The scripts for the latter analysis can be found in the GitHub repository (https://github.com/ravinpoudel/KEIO.git). For this analysis it is recommended to use an anaconda environment (anaconda.com) to run the KEIO.py and associated R scripts.

1.Run Illumina sequencing reads received as .fastq files through the FEBA program.
2.Convert TnPool.txt FEBA output file to .csv by changing the file extension.
3.Install Anaconda and create a conda environment with the following dependencies: VSEARCH, pybedtool, NMSLib, Biopython, and Pandas using the following code on the command prompt.
5. Inside the keio conda environment install the KEIO program from https://github.com/ravinpoudel/KEIO.git with the following commands
6. In the keio environment run KEIO.py with the following inputs: -- fastq: the forward Illumina sequencing file for the 384-well arrayed library. –upstreamFASTA: a .fasta file with the unique 16 BSup multiplexing primer sequences. –downstreamrcFASTA: a .fasta file with the unique 24 BSDown primers reverse complement sequences. This program returns the BSUp and BSDown primer pairs with their associated 20 bp random barcode as a results.csv file.
4. Download and install R (https://cran.r-project.org)
5. Create a results directory
6. Run the R script keio.R using the KEIO output.csv as the input. This program returns a .csv file output summarized.csv with the BSUp, BSDown and the 20 bp random barcode associated with the primers and their count statistics.
7. Download a general feature format file (.gff) and save it as a .csv file by changing the extension.
8. **Critical step**. Update the process.R script to tailor the gff.csv input to the desired organisms gff file.
9. Run the process.R script with the following inputs: the gff.csv file, the summarized.csv files and the Tnpool.csv FEBA output. This program outputs a final results.csv file. This file contains the 384 well plate 20 bp random barcode and associated statistics with the transposon location within the genome and genomic location annotations.

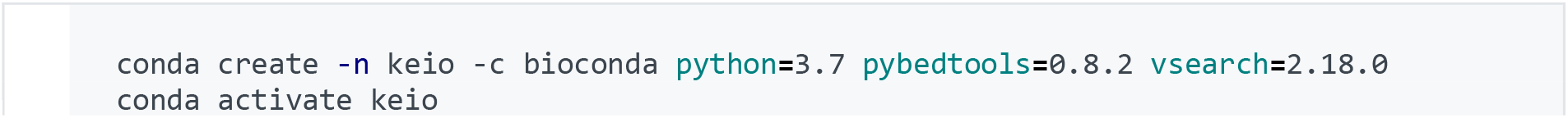

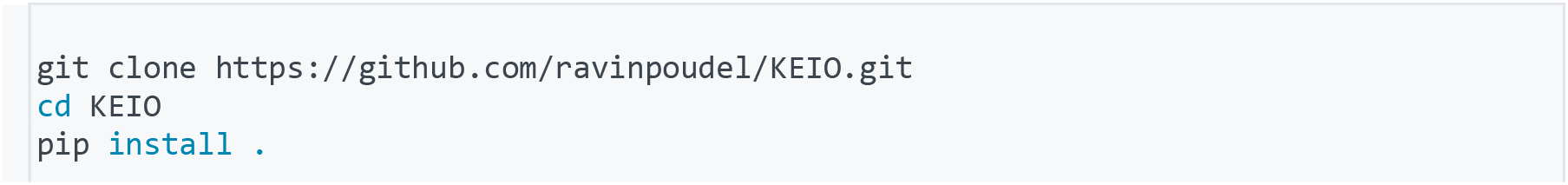

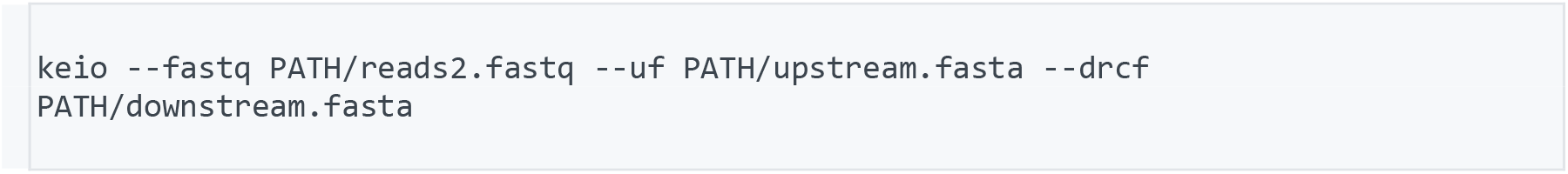

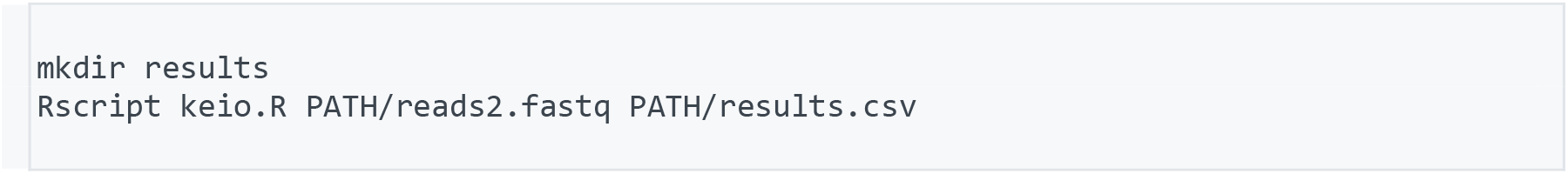

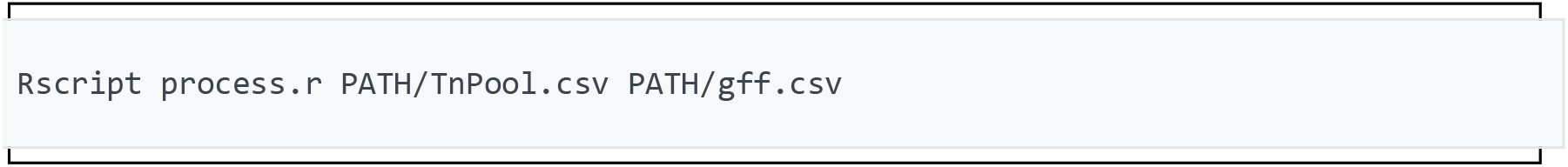

This pipeline will result in a final .csv file with the results for the 384-well plate random barcoded libraries. The final results.csv file will contain the well location for every mutant mapped with its corresponding 20 bp random barcode across all libraries. The results.csv file includes the 20 bp random barcode statistics as well as the transposon location for each mutant per well with its locus tag and other functional annotations.

## 5. REARRAYING

The arrayed 384-well libraries can be re-arrayed into 96- or 384-well plates to minimize the work required for genome wide screens. The Opentrons liquid handler is used to cherry-pick from the original plate into a new plate with other unique mutants.

1. Compile desired well locations from the arrayed 384-well libraries and allocate new well locations in destination source plates. Arrange this in a .csv format.
2. Prepare a .py script for the cherry-picking protocol, following the OT-2 Python Protocol API format.
3. Import .csv file into .py script. Perform a practice run of final .py as Python protocol on Opentrons liquid handler for further optimization.
4. Thaw glycerol 384-well source plate(s).
5. Aliquot 300 μL of ½ YTSS in deep 96-well plate(s). These will serve as destination microplates. Briefly spin down thawed source microplates to bring contents to the bottom of the wells.
6. Carefully remove foil on source microplates and cover with lids
7. Place source plate(s), destination plate(s), and tip rack (s) in designed slots of deck in Opentrons liquid handler.
8. Proceed to run cherry-picking protocol on Opentrons liquid handler.
9. Once cherry-picking is complete, prepare serial dilutions of source microplates in ½ YTSS at dilution factor necessary to obtain isolated colonies. Spot 10-20 μL of each series onto ½ YTSS kan.
10. Incubate spot plate (s) at 30° C for approximately 48 hours.
11. Inoculate isolated colonies into 384-well plates containing 80 μL of ½ YTSS kan. Incubate at 30° C for approximately 48 hours and replicate onto new 384-well plate containing 60 μL of ½ YTSS kan and store at –80° C.
12. Incubate replicated plate at 30° C for approximately 48 hours and save with 20 μL of 70% glycerol solution at –80° C. Follow previously described library arraying and plate mapping workflow (Sections 3 and 4) to confirm sequencing of rearrayed mutants.

## RESULTS

Using the methods described above, a total of four pooled transposon mutant libraries of *R. pomeroyi* were constructed to ensure a rich diversity of mutants. Random barcode transposon-site sequencing (RB-TnSeq) revealed 88.8% of all *R. pomeroyi* genes had a transposon insertion, with the transposon inserted in the same orientation as the gene 56.1% of the time. A total of 270,510 20-bp barcodes mapped to a transposon insertion in the genome. The pooled libraries were arrayed into 384-well plates inoculated with isolated colonies by an robotic colony picker, resulting in a total of 125 plates. The plate mapping protocol dubbed PlateSeq described here identified 27,448 of the 48,000 possible mutants in the arrayed plates (Table 1), while the remainder had barcodes that were not present in the TnSeq. This suggests that deeper sequencing in the TnSeq could lead to identification of more arrayed mutants. Regardless, the arrayed mutants represent 73.7% of all *R. pomeroyi* genes with a median of three mutants per gene.

**Table 1.**
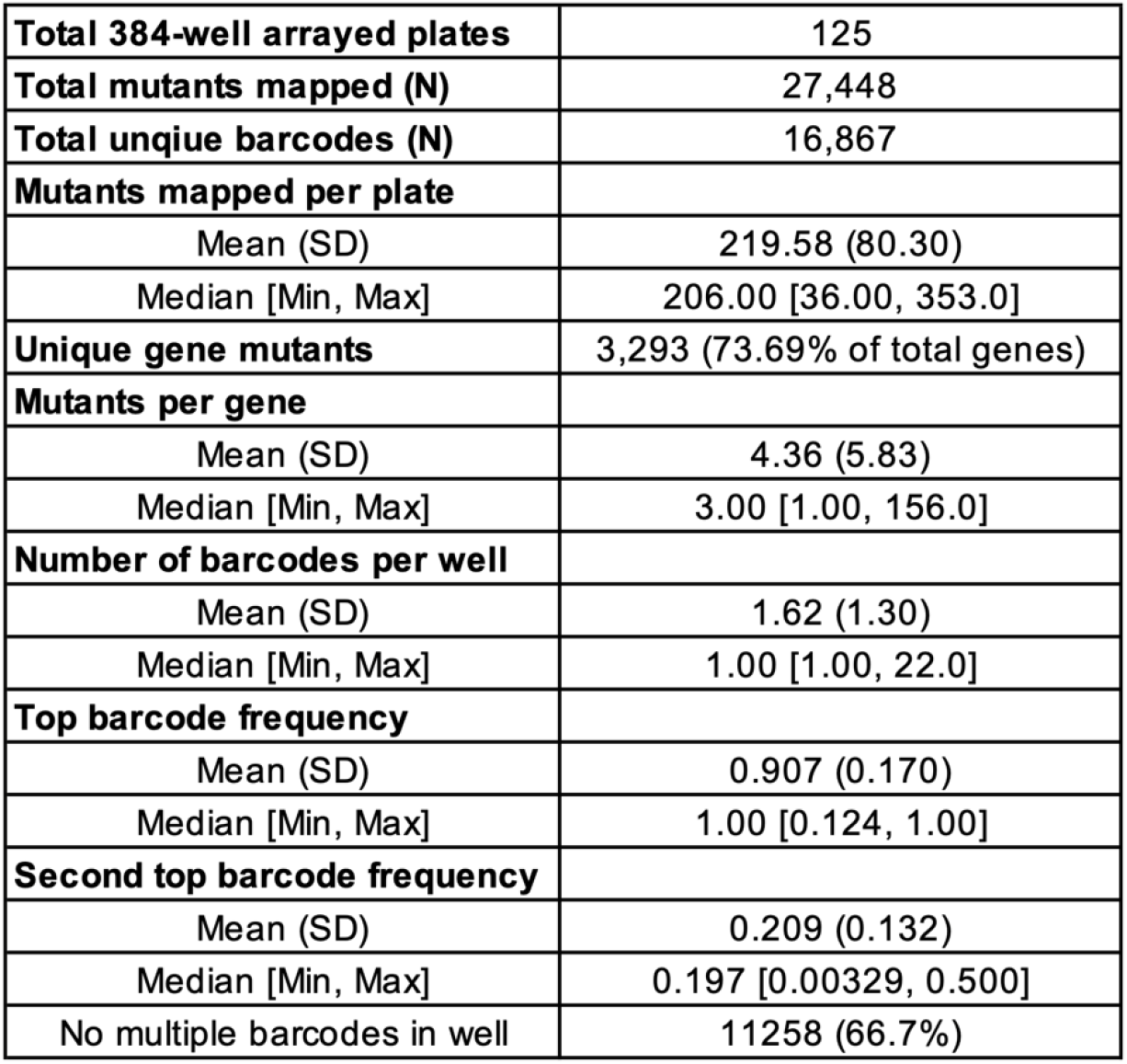
Statistics for the *R. pomeroyi* arrayed libraries.

PlateSeq analysis shows of the total arrayed library transposon mutants 90% map to the chromosome and 9.9% map to the megaplasmid of *R. pomeroyi* (Figure 2A). The arrayed libraries have a median of 1 barcode and an average of 1.6 barcodes sequenced per well (Table 1). In 66.7% of wells sequenced, only one unique barcode was mapped, indicating a single mutant in those wells (Table 1). In wells with multiple mapped barcodes the most abundant barcode accounts for an average of 91% of total reads for the well (Table 1). While the second most abundant barcode accounts for an average of 20% or less of the barcode reads of their well. Sanger sequencing of select arrayed mutants was used to verify that the barcode identified through NGS was in fact present in the well. Isolated colonies from 206 wells confirmed that 88.4% of wells matched the most abundant barcode from the NGS data (Figure 2B).

**Figure 2.**
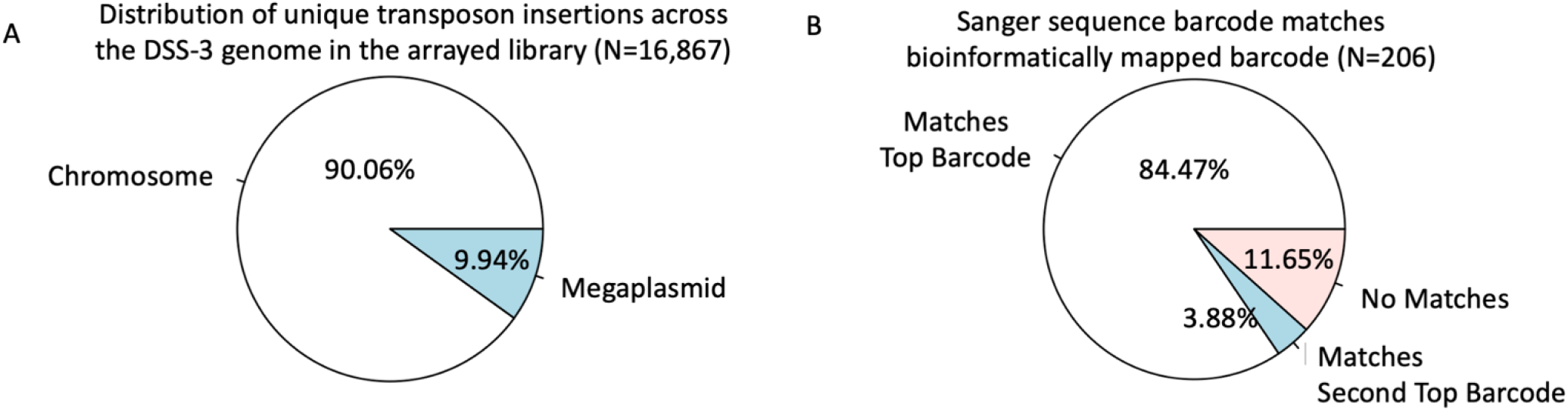
Overview of *R. pomeroyi* arrayed library results. (A) Distribution percentage of unique transposon insertions across the genome across arrayed library. (B) Arrayed library mutant barcodes were verified through Sanger sequencing for N=206 unique library mutants and compared to the computational mapping results.

Isolates from 8 of the remaining wells matched the second most abundant barcode found by NGS. These results confirm that there was in fact more than mutant present in many wells, though the correct mutant could still be isolated from a majority of wells.

## Acknowledgements

This work was supported by the USDA Agricultural Research Services (agreement #6066-21310-005-28-S), Simon’s Foundation, and National Science Foundation Center for Chemical Currencies on a Microbial Planet (OCE 2019589).

