## Supplementary Table 1 for "An Arrayed Transposon Library of *Ruegeria pomeroyi* DSS-3"

| BSDn1 | GAGTTCAGACGTGTGCTCTTCCGATCTCCATGTTGCCGGCCGTCGACCTGCAGCGTAC |
| --- | --- |
| BSDn2 | GAGTTCAGACGTGTGCTCTTCCGATCTACACCGGCCCGGCCGTCGACCTGCAGCGTAC |
| BSDn3 | GAGTTCAGACGTGTGCTCTTCCGATCTGTATCCTACCGGCCGTCGACCTGCAGCGTAC |
| BSDn4 | GAGTTCAGACGTGTGCTCTTCCGATCTGGATGAGCCCGGCCGTCGACCTGCAGCGTAC |
| BSDn5 | GAGTTCAGACGTGTGCTCTTCCGATCTGAGACTAGCCGGCCGTCGACCTGCAGCGTAC |
| BSDn6 | GAGTTCAGACGTGTGCTCTTCCGATCTGGCCTCTACCGGCCGTCGACCTGCAGCGTAC |
| BSDn7 | GAGTTCAGACGTGTGCTCTTCCGATCTCAATGATACCGGCCGTCGACCTGCAGCGTAC |
| BSDn8 | GAGTTCAGACGTGTGCTCTTCCGATCTCCTATCCACCGGCCGTCGACCTGCAGCGTAC |
| BSDn9 | GAGTTCAGACGTGTGCTCTTCCGATCTTTGATATACCGGCCGTCGACCTGCAGCGTAC |
| BSDn10 | GAGTTCAGACGTGTGCTCTTCCGATCTAGCGATATCCGGCCGTCGACCTGCAGCGTAC |
| BSDn11 | GAGTTCAGACGTGTGCTCTTCCGATCTCCTACAGTCCGGCCGTCGACCTGCAGCGTAC |
| BSDn12 | GAGTTCAGACGTGTGCTCTTCCGATCTATGACAGTCCGGCCGTCGACCTGCAGCGTAC |
| BSDn13 | GAGTTCAGACGTGTGCTCTTCCGATCTAGTGTACACCGGCCGTCGACCTGCAGCGTAC |
| BSDn14 | GAGTTCAGACGTGTGCTCTTCCGATCTCTGGCACGCCGGCCGTCGACCTGCAGCGTAC |
| BSDn15 | GAGTTCAGACGTGTGCTCTTCCGATCTCGCCTAACCCGGCCGTCGACCTGCAGCGTAC |
| BSDn16 | GAGTTCAGACGTGTGCTCTTCCGATCTCTCGTCGTCCGGCCGTCGACCTGCAGCGTAC |
| BSDn17 | GAGTTCAGACGTGTGCTCTTCCGATCTTACAGACACCGGCCGTCGACCTGCAGCGTAC |
| BSDn18 | GAGTTCAGACGTGTGCTCTTCCGATCTCAGTACCACCGGCCGTCGACCTGCAGCGTAC |
| BSDn19 | GAGTTCAGACGTGTGCTCTTCCGATCTAAGGTATCCCGGCCGTCGACCTGCAGCGTAC |
| BSDn20 | GAGTTCAGACGTGTGCTCTTCCGATCTAATTGAATCCGGCCGTCGACCTGCAGCGTAC |
| BSDn21 | GAGTTCAGACGTGTGCTCTTCCGATCTCGCAAGAGCCGGCCGTCGACCTGCAGCGTAC |
| BSDn22 | GAGTTCAGACGTGTGCTCTTCCGATCTCTCGATAACCGGCCGTCGACCTGCAGCGTAC |
| BSDn23 | GAGTTCAGACGTGTGCTCTTCCGATCTTTGTTCTCCCGGCCGTCGACCTGCAGCGTAC |
| BSDn24 | GAGTTCAGACGTGTGCTCTTCCGATCTTGACATCTCCGGCCGTCGACCTGCAGCGTAC |
| BSUp1 | CCTACACGACGCTCTTCCGATCTTCACTCACCGCCCTGCAGGGATGTCCACGAGGTCTC |
| BSup2 | CCTACACGACGCTCTTCCGATCTCGCCAGTACGCCCTGCAGGGATGTCCACGAGGTCTC |
| BSUp3 | CCTACACGACGCTCTTCCGATCTACTCAGGTCGCCCTGCAGGGATGTCCACGAGGTCTC |
| BSUp4 | CCTACACGACGCTCTTCCGATCTCGTAGCTTCGCCCTGCAGGGATGTCCACGAGGTCTC |
| BSUp5 | CCTACACGACGCTCTTCCGATCTCGCCTCAACGCCCTGCAGGGATGTCCACGAGGTCTC |
| BSUp6 | CCTACACGACGCTCTTCCGATCTAGTGATTCCGCCCTGCAGGGATGTCCACGAGGTCTC |
| BSUp7 | CCTACACGACGCTCTTCCGATCTATAAGAGGCGCCCTGCAGGGATGTCCACGAGGTCTC |
| BSUp8 | CCTACACGACGCTCTTCCGATCTAATGCCTTCGCCCTGCAGGGATGTCCACGAGGTCTC |
| BSUp9 | CCTACACGACGCTCTTCCGATCTTAGACTCCCGCCCTGCAGGGATGTCCACGAGGTCTC |
| BSUp10 | CCTACACGACGCTCTTCCGATCTACTATCTGCGCCCTGCAGGGATGTCCACGAGGTCTC |
| BSUp11 | CCTACACGACGCTCTTCCGATCTAGTTCGCACGCCCTGCAGGGATGTCCACGAGGTCTC |
| BSUp12 | CCTACACGACGCTCTTCCGATCTTCCGTACACGCCCTGCAGGGATGTCCACGAGGTCTC |
| BSUp13 | CCTACACGACGCTCTTCCGATCTGTGGATAGCGCCCTGCAGGGATGTCCACGAGGTCTC |
| BSUp14 | CCTACACGACGCTCTTCCGATCTAAGCTACGCGCCCTGCAGGGATGTCCACGAGGTCTC |
| BSUp15 | CCTACACGACGCTCTTCCGATCTCACACATCCGCCCTGCAGGGATGTCCACGAGGTCTC |
| BSUp16 | CCTACACGACGCTCTTCCGATCTATTCCTACCGCCCTGCAGGGATGTCCACGAGGTCTC |
